# Extreme Heat as the New Normal: A Methodological Roadmap for Behavior, Physiology, and Species Distributions

**DOI:** 10.64898/2026.02.25.707770

**Authors:** Diego Ellis-Soto, Daniel W. A. Noble, Pieter A. Arnold, Patrice Pottier, Alison J. Robey, Christina Prokopenko, Jeremy Cohen

## Abstract

A defining feature of climate change is the increasing frequency, intensity, and severity of extreme weather events. Among them, extreme heat is recognized as a critical driver of ecological and evolutionary change. Intense heat episodes can exceed physiological limits, alter animal movement, restructure geographic ranges, and increase extinction risk more than gradual changes to mean temperatures. Yet links between extreme heat events and organismal biology remain limited, in part because definitions and metrics are not standardized, and user-friendly workflows and guides are lacking for many biologists. We present a methodological roadmap, with reproducible code, for integrating extreme heat into studies of behavior, physiology, biophysical ecology, species distribution models (SDMs), and population dynamics. First, we provide standardized computational approaches to define and quantify extreme heat. Second, we fit species distribution models for California quail (*Callipepla californica*) that include an extreme heat metric and showcase improved predictions of habitat suitability, particularly at range edges. Third, we compute biophysical simulations to quantify exposure to thermal stress in Sleepy lizards (*Tiliqua rugosa*) across distinct macro- and microclimates. Finally, accounting for temporal autocorrelation in temperature profiles in population simulation models, we show that clustered heat extremes—missed by averages—can increase the risk of population collapse. As extreme heat events become more common, incorporating their dynamics is essential for understanding ecological and evolutionary change, designing experiments across species’ geographic ranges, and supporting conservation in a rapidly warming world. Together, these case studies illustrate a reproducible, organism-informed roadmap to integrate extreme heat into predictions of ecological impacts and inference across levels of biological organization under ongoing climate change.

## Introduction

### Climate Change: From mean temperature increases to extreme heat events

Climate change poses one of the most significant threats to global biodiversity by driving widespread shifts in species’ distributions (Chowdhury et al., 2025; Pecl et al., 2017) and population declines (Bellard et al., 2012; Urban, 2015). Temperature limits species’ geographic ranges because increased heat stress can limit animals’ ability to forage, reproduce, and survive (Huey & Kingsolver, 1989). Thus, understanding how thermal stress affects activity, reproduction, and survival is essential for predicting the redistribution and decline of biodiversity under climate change. Beyond gradual increases in mean temperature, a hallmark of anthropogenic climate change is the increase in severity, intensity and frequency of extreme weather events, such that extreme heat events are becoming the ‘new normal’ (Chen et al., 2025; Fischer et al., 2021; Intergovernmental Panel on Climate Change (IPCC), 2023; Stillman, 2019).

Unlike gradual shifts in mean temperatures, which some organisms may be able to buffer through behavior or acclimation (Huey et al., 2012), extreme heat events can rapidly push organisms beyond their physiological limits and tolerances (Buckley & Huey, 2016). This can pose a high risk of lethal overheating, population declines, or local extirpation (Bailey & van de Pol, 2016; Gunderson et al., 2017; Harris et al., 2018; Murali et al., 2023). Extreme heat can further exceed physiological tolerances, altering animal behavior and windows of activity for energy acquisition, ultimately limiting where taxa are able to survive, which constrains the boundaries of species distributions (Brouwers et al., 2013; Stillman, 2019; van den Bosch et al., 2025) (Maxwell et al., 2019).

Despite the biological relevance and importance of studying extreme heat events for conservation, incorporating such events into modeling biodiversity patterns, such as species distributions or organismal responses remains underexplored (but see (Cohen et al., 2025; Gunderson et al., 2017; Morán-Ordóñez et al., 2018; Pottier et al., 2025), and often limited to single extreme heat events opportunistically captured by monitoring (Bailey & van de Pol, 2016; Cohen et al., 2020; Pinsky et al., 2019; Welch et al., 2023). The last decade has seen computational advances to monitor and disentangle extreme heat anomalies in decadal climate forecasts (Stewart et al., 2021a), as well as in measuring and obtaining microclimate data at fine spatial and temporal scales through biophysical downscaling and advances in microclimate sensors (De Frenne et al., 2025; Kemppinen et al., 2024; Maclean et al., 2019; Maclean & Klinges, 2021; Meyer et al., 2023). Incorporating extreme heat into ecological studies necessitates frameworks that link climatic variability directly to organismal outcomes (Arnold et al., 2025; Buckley et al., 2025). For example, the thermal load sensitivity (TLS) mechanistic framework enables researchers to quantify how organisms accumulate, repair, and respond to thermal damage over time in ways that map directly onto physiological processes, making them important for understanding ecological responses to extreme heat (Arnold et al., 2025).

### Extreme heat impacts species distributions at large spatial scales

To understand how global change and thermal environments shape broad patterns of biodiversity, ecologists commonly use species distribution models (SDMs). These relate environmental conditions to species occurrence records to estimate species–environment relationships and predict suitable habitat (Ellis-Soto et al., 2021; Guisan & Thuiller, 2005). These models typically incorporate macroclimatic predictors such as mean climates and seasonality of temperature and precipitation. However, there has been growing interest in incorporating extreme weather events (Cohen et al., 2025; Maxwell et al., 2019; Stewart et al., 2021b). For example, recent work on modeling the distributions of 535 North American birds incorporating extreme heat, cold, and drought variables showed consistently smaller estimated range sizes and lower species richness patterns than when modeling with climate means alone, ultimately leading to more reliable biodiversity predictions (Cohen et al., 2025). Similarly, incorporating extreme weather variables in the modeling of 37 plant species in southeast Australia, delivered significant improvements in model performance and differences in spatial patterns were most pronounced at the predicted range margins (Stewart et al., 2021a). In both examples, species living in hotter, drier regions subject to high maximum temperatures were linked to improved prediction accuracy when extreme weather was incorporated into the models. Despite these advances, such metrics derived from coarser climatic conditions may fail to capture the fine-scale microclimatic conditions organisms directly experience locally (Kemppinen et al., 2024; Klinges et al., 2024).

### Microclimate ecology and hourly extreme temperatures

Factors such as topography, vegetation, and soil composition impact local variation in temperature, water availability, solar radiation, cloud cover, wind speed, and evaporation rates (Bramer et al., 2018). Thus, variation at the local microclimate scale can greatly exceed the thermal variability depicted in coarser climate products across entire continents (De Frenne et al., 2025; Kemppinen et al., 2024). Microclimates may be more or less stable or extreme than the projected mean temperature changes for the area, and microclimates can attenuate the impacts of extreme weather via microrefugia or as suitable stepping stones between core habitat (De Frenne et al., 2025; Ellis-Soto et al., 2023; Hannah et al., 2014; Scheffers et al., 2014). For example, small mammals in the Mojave Desert have greater resilience to extreme heat than birds because they can exploit burrow microrefugia, effectively reducing evaporative cooling demands and thermal exposure (Riddell et al., 2019, 2021). In a large-bodied heat-sensitive mammal, moose (*Alces alces*), wet soil was a key microhabitat feature used by animals to alleviate heat stress (Verzuh et al., 2023). Microclimates can manifest at the scale of a single leaf, with predictions of arthropod thermal performance and vulnerability to extreme heat greatly differing when considering air temperature, intact leaf temperature or leaves experiencing herbivory (Kerr et al., 2025; Pincebourde & Casas, 2019). Omitting microclimate information when modeling biodiversity patterns may further mischaracterize organismal exposure to heat stress and lead to inaccurate predictions of species extinction risk and potential range shifts under climate change (Maclean & Early, 2023; Soifer et al., 2025). In practice, however, coarser meso- and macroclimatic data remain more widely used, largely because global datasets from satellites and reanalysis products are widely available (e.g., WorldClim (Fick & Hijmans, 2017), even though they can be less directly relevant to organisms than microclimatic conditions. Bridging this mismatch often requires first downscaling or reconstructing microclimates from meso- and macroclimatic inputs (and local habitat structure), and then translating those microclimates into organism-specific exposure (Klinges et al., 2024, 2025). Addressing these limitations requires mechanistic frameworks capable of translating microclimatic variability into organism-specific thermal exposure.

### From species distributions to biophysical modeling under extreme heat

Biophysical ecology links organismal traits (e.g., thermal tolerance, metabolic demands) with environmental drivers (e.g., microclimate variation) and better inform predictions of extreme heat exposure on organisms and its impacts on energetic requirements (Kearney et al., 2009). This approach is now facilitated by mechanistic niche models, which integrate microclimates with heat-and water-balance models for both ectotherms and endotherms (Kearney, Briscoe, et al., 2021). These models enable biophysical, trait-based (e.g., critical and preferred body temperature limits, evaporative-cooling capacity and metabolic demands) forecasts to be rapidly deployed across broad landscapes, incorporating both meso- and microclimatic variables alongside taxon-specific physiological parameters (Riddell et al., 2023). By estimating the fine-scale environmental conditions organisms experience, these tools help understand how behavioral and physiological buffering can mitigate the impacts of extreme heat events (Kearney et al., 2009), and explain why similar environmental temperatures may result in either lethal or sub-lethal outcomes. Recent work leveraging field-validated simulations show that desert lizards pay a ‘cost-of-living squeeze’, with warming driving up maintenance costs while reducing foraging opportunities (Wild et al., 2025). Biophysical modeling of three bird species in arid environments closely matched observed evaporative water loss and resting metabolic rate during periods of heat of these taxa. This identified air temperatures between 30-40°C as a threshold for sustained sublethal fitness costs for these taxa, which exceeds viable thresholds for evaporative cooling (Conradie et al., 2023). Taken together, biophysical models offer a powerful tool to improve fine scale climate vulnerability assessments of organisms using mechanistically informed modelling approaches.

### Embedded thermal history and population responses to extreme heat

While projecting organismal responses to extreme heat is critically important, outcomes like persistence, extinction, and extirpation occur not at the level of individual organisms but rather at the level of whole populations. Many approaches to forecasting population vulnerability are correlative or rely on comparisons between organismal thermal tolerances and projected frequency distributions of temperatures through non-linear averaging (e.g., (Deutsch et al., 2008; Vasseur et al., 2014)). However, individual extreme heat events may catastrophically affect populations despite only marginally affecting average projected thermal distributions and population growth rates (Pottier et al., 2025). While the sharp increase in damage and death rates under stressfully hot temperatures is well appreciated in the organismal context (Jørgensen et al., 2019, 2021, 2022; Ørsted et al., 2022; Rezende et al., 2014, 2020), it has generally been overlooked at the population level. One promising approach to account for the disproportionately negative impacts of heat events is through population dynamical modeling. Embedding measurements of a population’s growth or decline rate under different environmental temperatures into population growth models allows for the tracking of population abundance through time (Duffy et al., 2022; Robey et al., 2025; Robey & Vasseur, 2024; Vasseur, 2020; Vasseur et al., 2025). Such approaches capture the potential risks of extinction driven by short-term, but highly stressful extreme heat events, which may drive dramatic short-term extinction risks even in populations whose long-term average growth rate (and subsequent projected abundance) is significantly positive (Robey et al., 2025; Robey & Vasseur, 2024).

### Aims

Extreme heat shapes life across Bartholomew’s scales of biological organization (Bartholomew, 1986), affecting physiology, behavior, populations, and geographic ranges across biogeographic, taxonomic, and spatio-temporal scales (Stillman, 2019). To address this, we provide a series of case studies with openly accessible code that links these scales and demonstrates how extreme heat can be quantified and integrated into ecological analyses. Our motivation is to equip researchers with practical tools to disentangle the impacts of extreme heat and to quantify, map, and assess its role across their own study systems. First, we showcase how to define, detect, and quantify extreme heat in space and time. Second, we demonstrate how to incorporate long-term climatological metrics of extreme heat and drought into species distribution models and provide a worked example for the California quail (*Callipepla californica*) to evaluate their influence on model predictions. Third, we illustrate how to leverage biophysical modeling for the Sleepy lizard (*Tiliqua rugosa*) in combination with estimates of extreme heat across its geographic range to highlight differences in daily activity budgets across different macro- and microclimates. Fourth, we extend these approaches to population-level simulations to reveal how the timing and clustering of extreme heat events can alter demographic trajectories and elevate extinction risk.

## Methods & Results

### (I) What is extreme heat and how do we measure it?

Defining extreme heat events accurately is key to understanding and quantifying their biological impacts. There are numerous climatological definitions of ‘heatwaves’ and ‘extreme heat’, implemented through dozens of different temperature indices (Karl et al., 1999; Perkins & Alexander, 2013; Robinson, 2001; Smith et al., 2013). Broadly, heatwaves are commonly defined as a period of consecutive days with exceptionally high temperatures relative to conditions for that region and time of year, often defined against a historical reference baseline (Robinson, 2001). Common approaches include percentile-based thresholds (e.g. temperatures exceeding the 90th or 95th percentile of historical daily maxima), for a minimum duration (e.g., 3 or 5 consecutive days), or indices that combine both daily maximum and minimum temperature (Perkins & Alexander, 2013). These definitional choices can substantially alter ecological inference (Xu et al., 2016). For example, shifting the reference baseline definition (e.g., from 1979-2017 to 2000–2024) will change percentile thresholds, the number, timing, and intensity of extreme heat events. Furthermore, different percentile thresholds and the necessary number of consecutive days across such thresholds will further dramatically alter estimates of how often organisms are exposed to extreme heat. Similarly, relying on daily means or maxima can miss sub-daily thermal peaks, particularly when organisms experience short, intense periods of overheating.

Historically, biodiversity research has relied heavily on long-term climatological averages, exemplified by widely used products such as BIOCLIM variables from global datasets like WorldClim and CHELSA (Fick & Hijmans, 2017; Karger et al., 2017) jointly cited in well over 20,000 publications. These variables, over periods exceeding 30 years, describe broad-scale seasonal patterns but fail to capture short-term extreme events that can lead to acute physiological stress or increased mortality and population declines. New climatological products and computational advances offer more customized approaches. Data infrastructures such as NASA’s Distributed Active Archive Centers, the Copernicus Data Space Ecosystem, and national climate archives now serve daily to sub-daily gridded temperature fields at increasingly fine spatial resolutions. Open source packages in the R Environment for statistical computing (R Core Team, 2024), such as *daymetR*, *heatwaveR*, now allow retrieving time series of temperature for specific locations and periods (Hufkens et al., 2018; Schlegel & Smit, 2018). These allow users to calculate custom indices (e.g., exceedance of thermal thresholds, cumulative heat load), and derive event-level metrics such as onset, duration, and peak intensity (Crego et al., 2021; Dodge et al., 2013; Li et al., 2021). Geospatial platforms like Google Earth Engine and Microsoft Planetary Computer (https://earthengine.google.com/, https://planetarycomputer.microsoft.com/), further enhance the accessibility and scalability of these analyses, offering biologists flexible and powerful interfaces to customize extreme heat predictor variables into their ecological research rather than using pre-aggregated climatologies (Gorelick et al., 2017). High-resolution products like ERA-5 with the associated R package *mcera5* allow obtaining hourly temperature estimates to quantify intra-day thermal stress during extreme heat events (Klinges et al., 2022), complementing traditional measures such as daily maximum temperatures. Taken together, these methodologies allow quantifying temperature anomalies—defined as deviations from historical daily means or other baselines—and provide a flexible way to detect and compare extreme events across locations, years, and climate scenarios. These can be computed at daily or sub-daily resolution to match organismal time scales.

In our case study in Figure 1, we illustrate how to quantify and identify periods of extreme heat using custom thresholds for a particular point of interest (Latitude 48.3, Longitude -117.5, Washington State, United States of America). We first used *daymetR* to retrieve daily maximum temperature (Tmax) from the Daymet reanalysis product at 1 km^2^ scale for both the historical baseline period (1991–2020) and a focal year (2021) at a specified location. We then used *heatwaveR* to identify and characterize extreme heat events in the 2021 time series, extracting metrics such as event duration, maximum intensity, and cumulative thermal stress. In doing so, we quantified extreme air temperatures experienced by animals at a site of interest without considering behavioral buffering, landscape features and microclimates (Figure 1). This workflow provides a reproducible way to quantify extreme air temperatures for organisms experiencing the June–July 2021 heat dome event, one of the most intense extreme events in the western United States in the last 1000 years (Thompson et al., 2022). A limitation of our case study is that *daymetR* is only available for the United States, and alternative methodologies could leverage MODIS Land Surface Temperature, which is available globally at the daily scale for day and nighttime temperatures.

**Figure 1:**
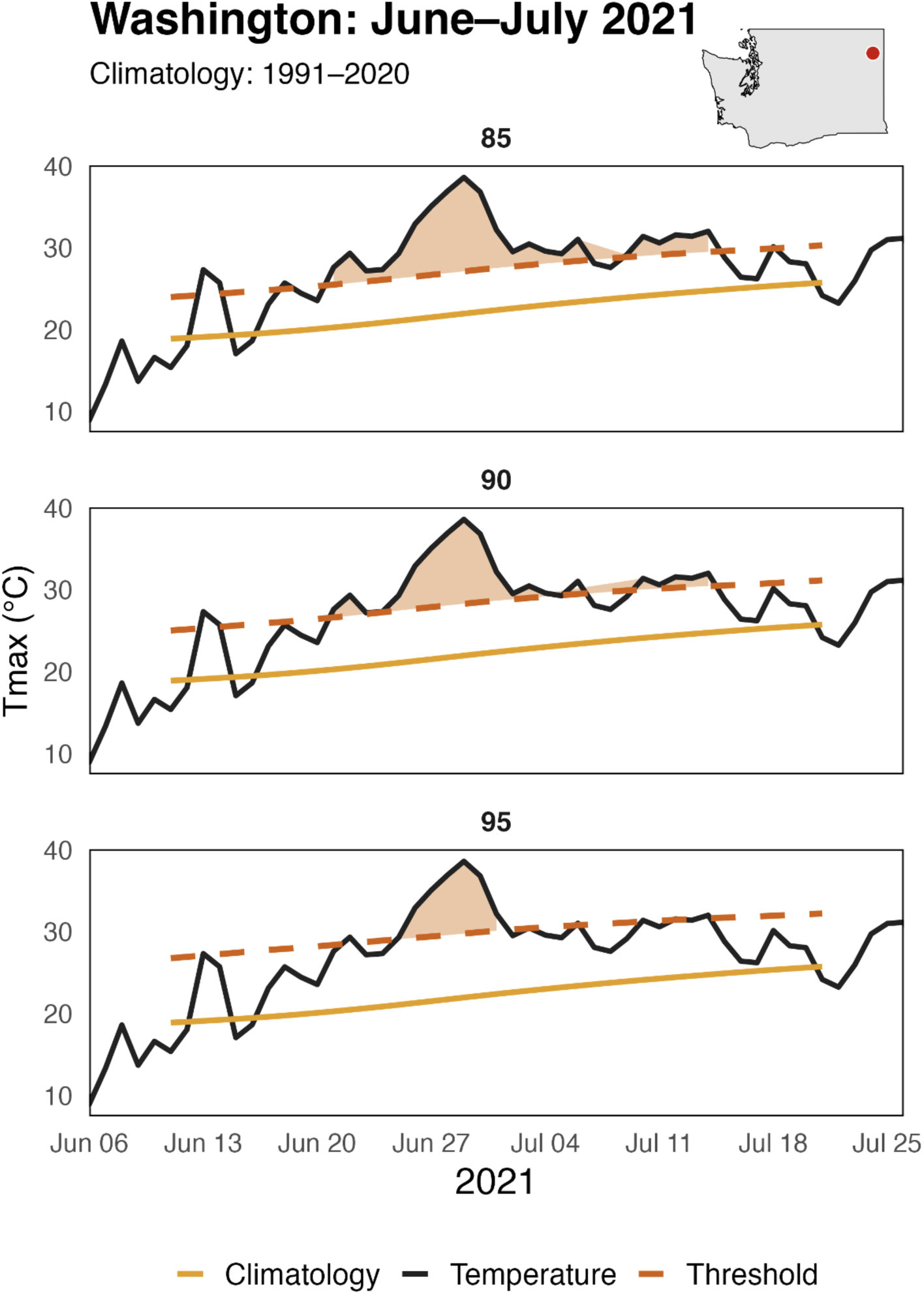
Identification of an extreme heat event with its respective duration, magnitude and time period in Washington State, United States. Daily maximum air temperature (Tmax) time series at 1-km spatial resolution. The dashed red line represents a defined extreme-heat threshold (here 85th, 90th and 95th percentile, respectively across top, middle and bottom panels). Shaded light red regions indicate periods of extreme heat, defined as intervals in which observed Tmax exceeds the percentile-based threshold for a minimum number of consecutive days (here ≥5 days), which is a user-defined customizable parameter. The orange line shows the daily climatological mean Tmax, calculated for each day of the year as the average across the 1991–2020 baseline period. This highlights the importance of jointly considering heat intensity, threshold to identify extreme heat, and duration when linking environmental temperatures to organismal stress and biological outcomes. Days exceeding the threshold but not meeting the minimum consecutive-day requirement are not classified as extreme heat events; this duration threshold is user-defined and adjustable. We provide necessary code to map and visualize extreme heat across multiple thresholds across the United States of America in the associated GitHub repository.

### (II) Incorporating extreme weather into species distribution modeling

To illustrate how extreme heat events shape species distributions, we integrated extreme weather risk maps into SDMs for the California quail (*Callipepla californica*), a species known to be sensitive to extreme heat (Perry et al., 2022). Using observational data from 222,971 eBird checklists, we constructed three random forest species distribution models across the United States, incorporating climatic means, climatic variability (seasonality), and extreme heat and drought risk layers derived from high-resolution (1 km²) temperature extremes datasets (https://envidat.ch/)(La Sorte et al., 2021). For climatic means, we incorporated mean annual temperature and mean annual precipitation (bioclim variables 1 and 12), while climatic variability was represented by temperature seasonality and precipitation seasonality (Bioclim 4 and 15, respectively) from Chelsa (Karger et al., 2017). We compared these three predictions, building upon previous research when assessing the influence of extreme weather on resident avian biodiversity across the United States (Cohen et al. 2025). These extreme weather layers captured the duration and intensity of both extreme heat events (EHE) and drought conditions, assessed via the Standardized Precipitation Evapotranspiration Index (SPEI), providing insights into the impacts of episodic climatic stress on species occurrence. We used the *raster* (Hijmans et al., 2015) and *sf* (Pebesma, 2018) packages in R for geoprocessing and the *randomForest* package for modeling (Liaw & Wiener, 2002).

Our results indicate substantial improvements in predictive accuracy when incorporating extreme weather into SDMs, particularly in identifying unsuitable habitats where frequent temperature extremes limit species presence (Figure 2). Specifically, models accounting for extreme weather events demonstrated notably reduced range predictions, particularly at distributional edges where physiological limits are approached (Figure 2). For instance, predictions without extremes overestimated the quail’s distribution into interior regions characterized by larger temperature fluctuations and underestimated their presence in milder coastal regions such as Baja California. Partial dependence analyses further revealed a clear negative relationship between the probability of occurrence and both heat and drought extremes. Overall, model specificity, or the ability to correctly predict absence observations improved from 0.835 to 0.851 with the inclusion of extreme weather variables (alongside a model AUC improvement of 0.78 to 0.80), highlighting that extreme weather acts as an important but often omitted environmental filter influencing the persistence of organism. In fact, our model-estimated area of occurrence declined 9% (1.087 to 0.989 million km2) with the inclusion of extreme weather, suggesting that models lacking this filter could overpredict species ranges. More generally, this example illustrates the potential of incorporating extreme climatic events for ecological forecasting.

**Figure 2:**
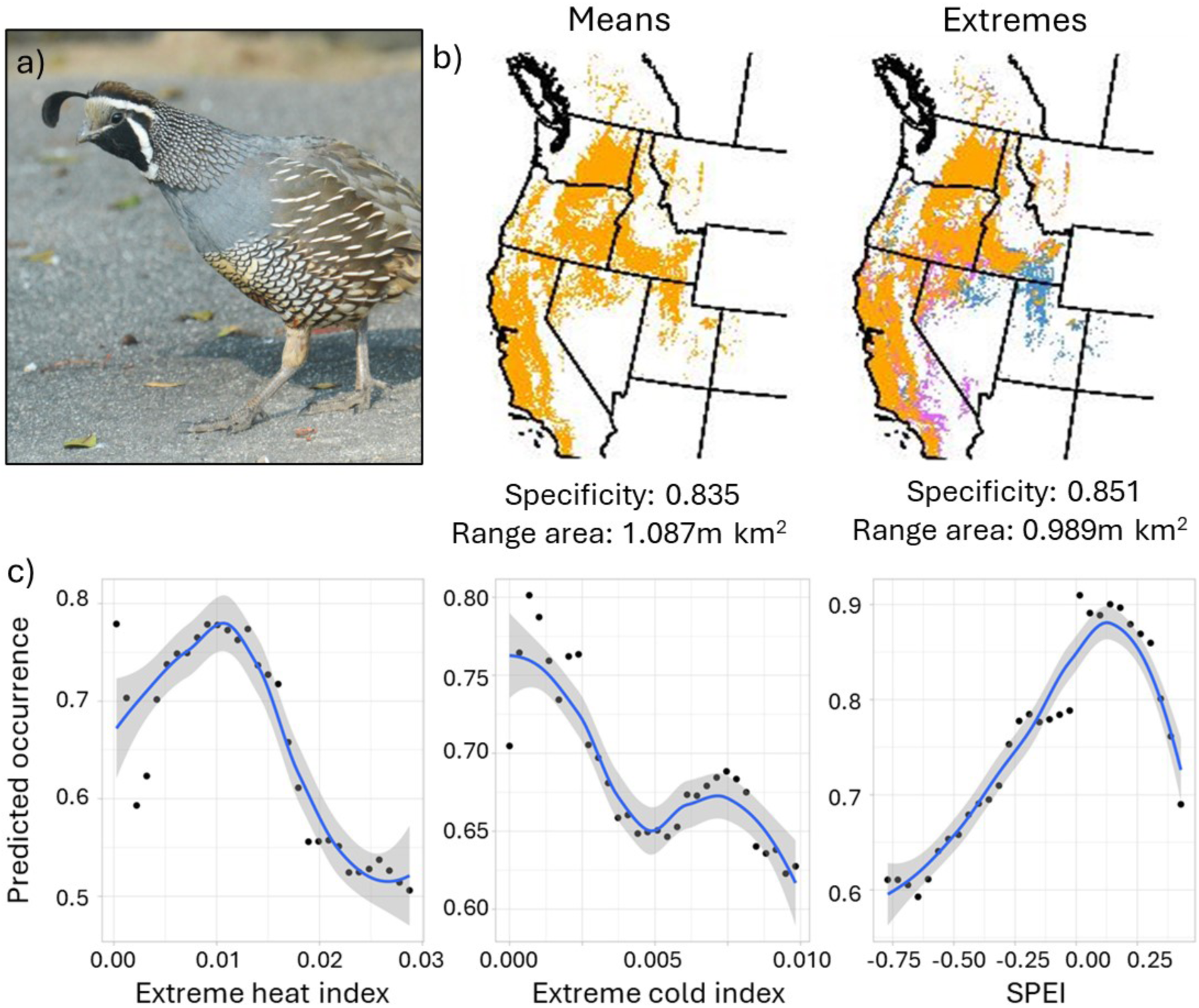
Bird distributions in the context of extreme weather. Predicted breeding range of California quail (*Callipepla californica*) across models including b) only climate means (left) or climate variability and extreme weather risk (right). Orange areas represent predicted suitable areas in both models and blue areas are not suitable in the extreme weather model despite suitability in models with only climatic means, while purple areas become suitable only when climate extremes are included. The known breeding range of this species does not extend as far into eastern Utah, Idaho and Nevada as suggested by the model containing only climatic means, but does include southeastern California. In (c), partial dependence plots from the extreme weather model convey the estimated relationship between the likelihood of occurrence and (left to right) the extreme heat index (high risk represented by high values), extreme cold index (high values), and Standardized Precipitation Evapotranspiration Index (SPEI, negative values).

### (III) Biophysical ecology and microclimates: Mechanistic niche modeling and microclimates

Broad climatic variables ignore the microclimatic heterogeneity that organisms can exploit to mitigate exposure to extreme heat. Microclimatic conditions can be estimated for a given location and integrated with biophysical models that incorporate heat/water budgets and behavioral responses to calculate realised organismal temperatures (Briscoe et al., 2023; Kearney, Briscoe, et al., 2021; Kearney, Porter, et al., 2021). These approaches can provide more realistic predictions of extreme heat impacts because they estimate what conditions organisms are actually experiencing during periods of thermal extremes. As an example of how such approaches can provide more nuanced predictions, we estimated the realized body temperatures of an ectothermic organism (sleepy lizard, *Tiliqua rugosa*) that is broadly distributed across southern Australia in temperate and semiarid ecosystems. Here, we use species-specific parameters that have been ground-truthed in field studies (Kearney et al., 2018) to fit the ectotherm model with *NicheMapR* (Kearney, Briscoe, et al., 2021; Kearney, Porter, et al., 2021). This approach integrates preferred body temperature, thermal tolerance limits, and behavioral thermoregulation strategies (e.g., shade seeking and burrowing) to attempt to maintain a preferred body temperature of 33.5°C with the microclimates that are available to the organism.

Here we contrasted two locations (coastal and inland) that differ in their number of extreme heat days (% of days > TX90p defined as the proportion of days exceeding the 90th percentile threshold for daily maximum temperature of a baseline period (Zhang et al., 2005); Figure 3). We then simulated the microclimate and body temperature of *T. rugosa* at each location during a window of 10 hot days in January (Austral mid-summer). Then, we calculated the mean thermal safety margin (TSM) on an hourly basis between the critical thermal maximum (CTmax) of the organism (42°C) and daytime (06:00–19:00) body temperatures. We compared these predictions to those based on the typical approach using mesoclimate data: calculations of TSM using CTmax and maximum annual air temperature from Worldclim (‘Bio5’) at the same locations, which is more coarse and does not account for biophysical principles and behavior.

**Figure 3:**
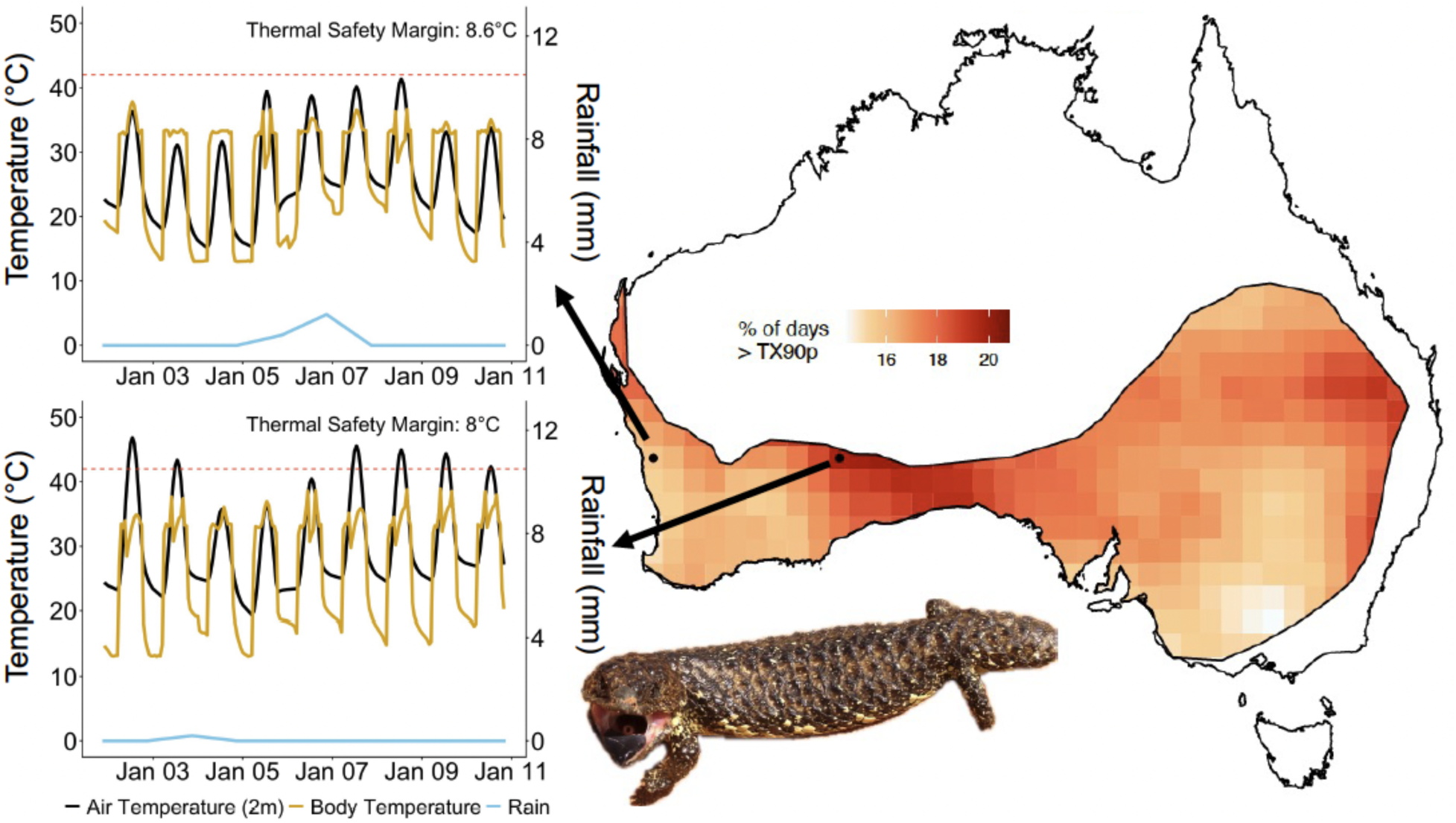
Air temperatures across different microclimates as well as realised body temperatures predicted for two populations of Sleepy lizards across its geographic range. Rainfall is also provided at these locations as this impacts microclimate and organismal body temperatures. Lizards were allowed to behaviorally thermoregulate exploiting microhabitat heterogeneity to try and maintain body temperatures around Tpref = ∼33°C. Thermal Safety Margins between realised body temperature and upper thermal limit (CTmax = 42°C) are also presented demonstrating how behavioral plasticity can mitigate extreme heat events.

At the coastal location, air temperatures were generally lower and body temperatures exceeded the preferred temperature less frequently than the inland location (Figure 3). The median thermal safety margin was similar between locations due to behavioral thermoregulation. Calculations of TSM using WorldClim showed TSMs were between 0.3-2.0°C lower for each site, and that the sites were more different to each other compared to the biophysical models that incorporate behavior (coastal: 6.6°C vs 8.6°C; inland: 7.7°C vs 8.0°C). This difference (albeit small) indicates that behavioral plasticity mitigates many of the potential increases in body temperature at the inland location, despite the higher air temperatures. This is reinforced by observations that running the ectotherm model without permitting burrowing (i.e., only shade-seeking behavior), leads to CTmax being exceeded on 7 out of 10 days at the inland location. We expect such behaviorally-mediated differences to be even greater for species living in more complex habitats. Nonetheless, this lays the groundwork for further understanding how these animals may behave in response to microclimates and adjust energetic costs to thermal extremes. This integration of high-resolution microclimate simulations with biophysical and physiological modeling provides a powerful framework for mechanistically predicting species’ responses to increasingly frequent and intense periods of extreme temperature.

### (IV) Embedded thermal history and population responses to heatwaves

To illustrate the effect of extreme heat on population outcomes, we embedded thermal sensitivity into a logistic model of population growth. The change in population size *N* at any given time *t* can thus be calculated as *dN*/*dt* = *N* [*r*(*T*) − *αN*], where *α* represents the strength of density dependence and *r*(*T*) is the instantaneous population growth rate at temperature *T* (Long et al., 2007; Mallet, 2012; Vasseur, 2020). Population growth rate is positive over the range of temperatures where birth rate outpaces death rate (and negative where death outpaces birth). Most organisms exhibit left-skewed population growth rate thermal performance curves (TPCs; (Kontopoulos et al., 2024; Rezende & Bozinovic, 2019); e.g., Figure 4a), where growth rate gradually increases to a peak at the thermal optimum, then rapidly declines with further warming (Angilletta Jr., 2009). Incorporating TPCs into population dynamical modeling allows us to track variations in abundance (and subsequent persistence or extinction outcomes) according to the temperature-dependent rates of population growth and decline for explicit temperature time series (Duffy et al., 2022; Robey & Vasseur, 2024; Vasseur, 2020). Previous approaches assessing risk based on averaging implicitly assume temperatures occur in a random order through time (i.e., no temporal autocorrelation). However, on the timescale relevant to population growth rates, temperatures typically exhibit positive autocorrelation (where similar temperatures cluster together) (Vasseur & Yodzis, 2004). For a given thermal distribution, the likelihood of longer, more proximate heat events thus increases with positive autocorrelation, leading extinction risks to grow beyond what averaging approaches predict.

**Figure 4:**
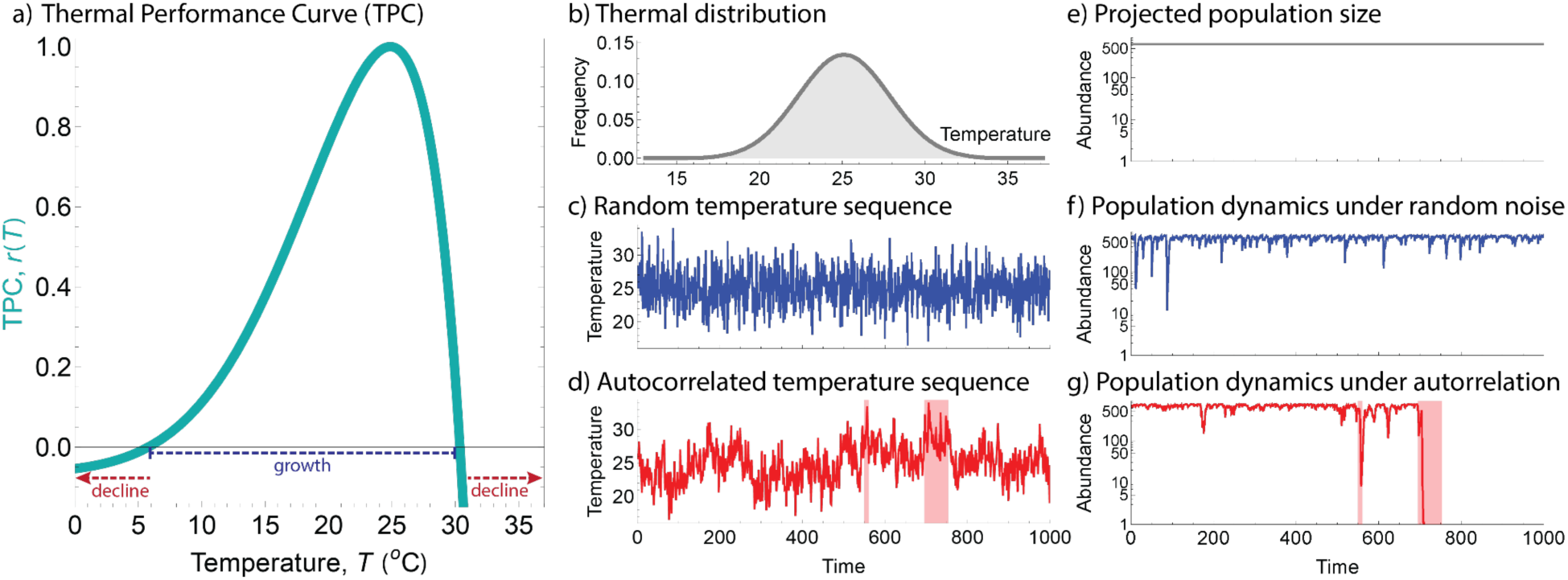
Thermal performance curves and temperature variability influence population dynamics. (a) Normalized thermal performance curve (TPC) of a population’s growth rate *r* at each temperature *T*, with an optimum at 25°C 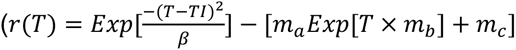, with *TI* = 29, *β* = 180, *m_a_* = 5 × 10^-6^, *m_b_* = 0.4, *m_c_* = 0.05). (b) Environmental temperature distribution, normally distributed with a mean of 25°C and standard deviation of 2.75°C. (c) Example randomly-ordered sequence of the thermal distribution (spectral exponent *γ* = 0); such random ordering is the implicit assumption of averaging approaches that do not account for the sequence of events. (d) Example positively autocorrelated sequence of the thermal distribution (*γ* = 1), with clusters of stressfully hot temperatures highlighted in red. (e-g) Projected population outcomes using each assumptions in (b-d; shown dynamics were run with *α* = 0.001); although logistically growing populations usually have abundances close to (e) the average size forecasted using only the thermal distribution (*N* = 632), populations (f) fluctuate more when dynamics are included and (g) risk extinction under more realistic levels of temporal autocorrelation. Adapted from Robey & Vasseur (2026).

Embedding thermal history into population modeling demonstrated the disproportionate effect of such heat events. For example, we showed an environmental temperature distribution (Figure 4b) that appeared well-matched with a population’s growth rate TPC (Figure 4a); although this thermal distribution included stressful temperatures (i.e., where *r*(*T*) < 0 and the population shrinks), the average growth rate was positive (Figure 4e) and persistence was expected. While embedding an explicit (albeit random) series of temperature events (Figure 4c) into population dynamical models induced some fluctuations (Figure 4f), the population retained a high abundance well-predicted by the averaging approach (Figure 4e). However, when the same thermal distribution exhibited positive autocorrelation (Figure 4d), heatwaves were generated (red shading) which, when filtered through the population dynamics, led to an extinction (Figure 4g). Accounting for the temporal clustering of heat events by explicitly including the ordering of forecasted temperatures into risk projections is key to understanding population resilience to stressful heat events (Ma et al., 2018; Robey & Vasseur, 2024).

## Discussion

Our case studies provide a reproducible roadmap for linking extreme heat to organismal behavior and physiology, biogeography, and population outcomes. Through species distribution models for California quail, we show that incorporating extreme heat and drought metrics substantially improves forecasts by reducing the overestimation of range boundaries that arises when models rely solely on climatic means and seasonality (Figure 2). Our biophysical simulations for Sleepy lizards further reveal how microclimate heterogeneity and behavioral thermoregulation modulate organismal exposure to thermal stress (Figure 3), underscoring the importance of mechanistic frameworks that incorporate fine-scale environmental variation. Finally, population dynamic simulations illustrate that the timing and clustering of heat extremes, not only their long-term averages, can fundamentally alter demographic trajectories and elevate extinction risk (Figure 4).

Moving forward, the increasing availability of high-resolution climatic datasets and advances in microclimate modeling create opportunities to extend classical climatological definitions of heatwaves toward organism-centric, mechanistic frameworks. Future studies quantifying extreme heat could more clearly state and justify the selected thresholds or indices, acknowledge their limitations, and evaluate the robustness of ecological inferences across alternatives given how such choices influence estimates of exposure and the biological conclusions drawn from them (Figure 1). Frameworks such as the thermal load sensitivity (TLS) approach offer a complementary approach by integrating heat magnitude and duration with physiological processes of damage and repair, extending classical thermal-death-time models (Arnold et al., 2025). Because TLS links environmental temperature regimes directly to organismal performance, survival, and demographic outcomes, it enables biologically informed thresholds to be derived from damage accumulation or key thermal traits (e.g., CTmax). This allows both cumulative and episodic thermal stress to be quantified consistently across taxa and event types (from brief thermal spikes to prolonged warm periods) even when conditions do not meet formal climatological definitions of extreme heat or a heatwave.

With the increasing availability of wildlife mortality records in near real time (Ellis-Soto et al., 2025), there is ample opportunity to assess whether extreme heat events coincide with reports of focal species deaths. Such records can strengthen associations between organismal stress and demographic consequences. As fine-scale climatic datasets and computational tools increasingly allow detailed characterization of thermal exposure, pairing these advances with integrative frameworks that account for mortality or other consequences will facilitate more biologically grounded assessments of extreme heat in ecological and evolutionary research. Adopting such integrative approaches will be essential for predicting biological responses and informing conservation strategies as extreme heat becomes an increasingly dominant feature of the Anthropocene.

## Code availability statements

We provide the code generated for our manuscript analysis on https://github.com/diego-ellis-soto/Extreme_heat_SICB

## Acknowledgements

We are grateful for input from Owen Atkin and Michael R. Kearney. DES acknowledges funding support from the David H. Smith Research Fellowship and the Presidential Postdoctoral Program of the University of California.

## Author contributions

DES contributed to conceptualization and led manuscript writing with substantial contribution from all co-authors. DES, DWAN, JC, PAA and AJR designed methodology, performed analyses and generated figures. All co-authors contributed to reviewing and editing the manuscript draft.

